# A computational mechanism for seeing dynamic deformation

**DOI:** 10.1101/682336

**Authors:** Takahiro Kawabe, Masataka Sawayama

**Author notes:** Corresponding author 3-1, Morinosato Wakamiya, Atsugi, Kanagawa, 243-0198, Japan.

## Abstract

Human observers perceptually discriminate the dynamic deformation of materials in the real world. However, the psychophysical and neural mechanisms responsible for the perception of dynamic deformation have not been fully elucidated. By using a deforming bar as the stimulus, we showed that the spatial frequency of deformation was a critical determinant of deformation perception. Simulating the response of direction-selective units (i.e., MT pattern motion cells) to stimuli, we found that the perception of dynamic deformation was well explained by assuming a higher-order mechanism monitoring the spatial pattern of direction responses. Our model with the higher-order mechanism also successfully explained the appearance of a visual illusion wherein a static bar apparently deforms against a tilted drifting grating. In particular, it was the lower spatial frequencies in this pattern that strongly contributed to the deformation perception. Finally, by manipulating the luminance of the static bar, we observed that the mechanism for the illusory deformation was more sensitive to luminance than contrast cues.

**Significance Statement:** From the psychophysical and computational points of view, the present study tried to answer the question, “how do human observers see deformation?”. In the psychophysical experiment, we used a clip wherein a bar dynamically deformed. We also tested the illusory deformation of a bar, which was caused by tilted drifting grating, because it was unclear whether the illusory deformation could be described by our model. In the computational analysis, in order to explain psychophysical data for deformation perception, it was necessary to assume an additional unit monitoring the spatial pattern of direction responses of MT cells that were sensitive to local image motion.

## Introduction

Materials in the real world are often non-rigid. The material non-rigidity dynamically produces the deformations of contours and textures in the retinal images. The dynamic deformation of retinal images is a rich source of visual information allowing the visual system to assess material properties in the real world. For example, from the dynamic deformation of retinal images, human observers can recognize transparent liquid (Kawabe, Maruya, & Nishida, 2015), transparent gas (Kawabe & Kogovšek, 2017), the elasticity and/or stiffness of materials (Masuda et al., 2013, 2016; Paulun et al., 2017; Schmidt et al., 2017), the stiffness of fabrics (Bi & Xiao, 2016; Bi et al., 2018; Bi et al., 2019), biological motion (Blake & Shiffrar, 2007; Johansson, 1973; Kawabe, 2017), and more.

Importantly, however, the visual mechanism for detecting dynamic deformation itself has not been thoroughly examined. Previous studies have reported that observers reported dynamic deformation when local motion integration did not provide evidence for rigid motion (Nakayama & Silverman, 1988a, 1988b) at a layer-represented level (Weiss & Adelson, 2000). Although successfully addressing the stimulus condition in which rigid motion perception was violated, these previous studies have not mentioned the conditions to cause the perception of dynamic deformation. Although some studies have proposed the computational model for the detection of two-dimensional motion patterns containing shearing and/or rotating motion (Sachtler & Zaidi, 1995; Zhang, Sereno, & Sereno, 1993), the model is not directly considered as the explanation of deformation perception because, as shown in a previous study (Nakayama & Silverman, 1988a), a shearing motion pattern does not always cause deformation perception. Nakayama and Silverman (1988a) showed that dynamic contour deformation with higher deformation frequency did not produce deformation perception but caused the rigid movement of wavy patterns, indicating that the detection of the shearing motion itself does not always lead to shearing deformation perception. No previous computational model exactly accounts for the dependency of deformation perception on the spatial frequency of deformation. Hence, additional examinations are necessary to fully understand mechanisms for deformation perception in human observers. Moreover, it remained unclear what visual information could be effective in generating the representation of dynamic deformation. Jain and Zaidi (2011) have shown that motion is important information for discerning the shape of non-rigidly deforming objects. The present study thus focuses on how motion information contributes to the perception of dynamic deformation.

The purpose of this study was to psychophysically and computationally specify the mechanism that underlies the perception of dynamic deformation. In Experiment 1, using stimuli with a physically deforming bar we show that the spatial frequency of deformation is an important factor to phenomenally determine deformation perception. By simulating both spatiotemporal energy responses at the V1 level and the responses of direction-selective units [i.e., MT pattern motion cells (Perrone & Krauzlis, 2008; Perrone & Krauzlis, 2014; Simoncelli & Heeger, 1998)], we show that not spatiotemporal motion energy but the spatial pattern of the responses of the direction-selective units consistently explains observers’ reports for the perception of dynamic deformation. In Experiment 2, we first examine a visual illusion in which a static bar with solid edges apparently deforms against a slightly tilted drifting grating (See Figures 4a and 4b). We report that both the orientation and spatial frequency of the background grating are critical to the illusory perception of deformation. From the observation, it is plausible to assume that Moiré patterns (Oster, 1965; Spillmann, 1993; Wade, 2007), which are generated between the bar’s edge and background grating, produce motion signals that are related to the apparent deformation. However, it is still unclear whether the dependence of the deformation appearance on both orientation and spatial frequency can be explained by the spatial pattern of the responses of direction-selective units. We again analyze the spatial pattern of the direction-selective unit responses to the illusion display and examine whether they again predict the observers’ report of deformation in the illusory deformation. In Experiment 3, we show that the perception of illusory dynamic deformation is attenuated when the average luminance of background grating is equivalent to the luminance of a bar. We then discuss how the perception of dynamic deformation is determined on the basis of the output of a high-level mechanism monitoring the spatial patterns of responses of direction-selective units.

**Figure 1.**
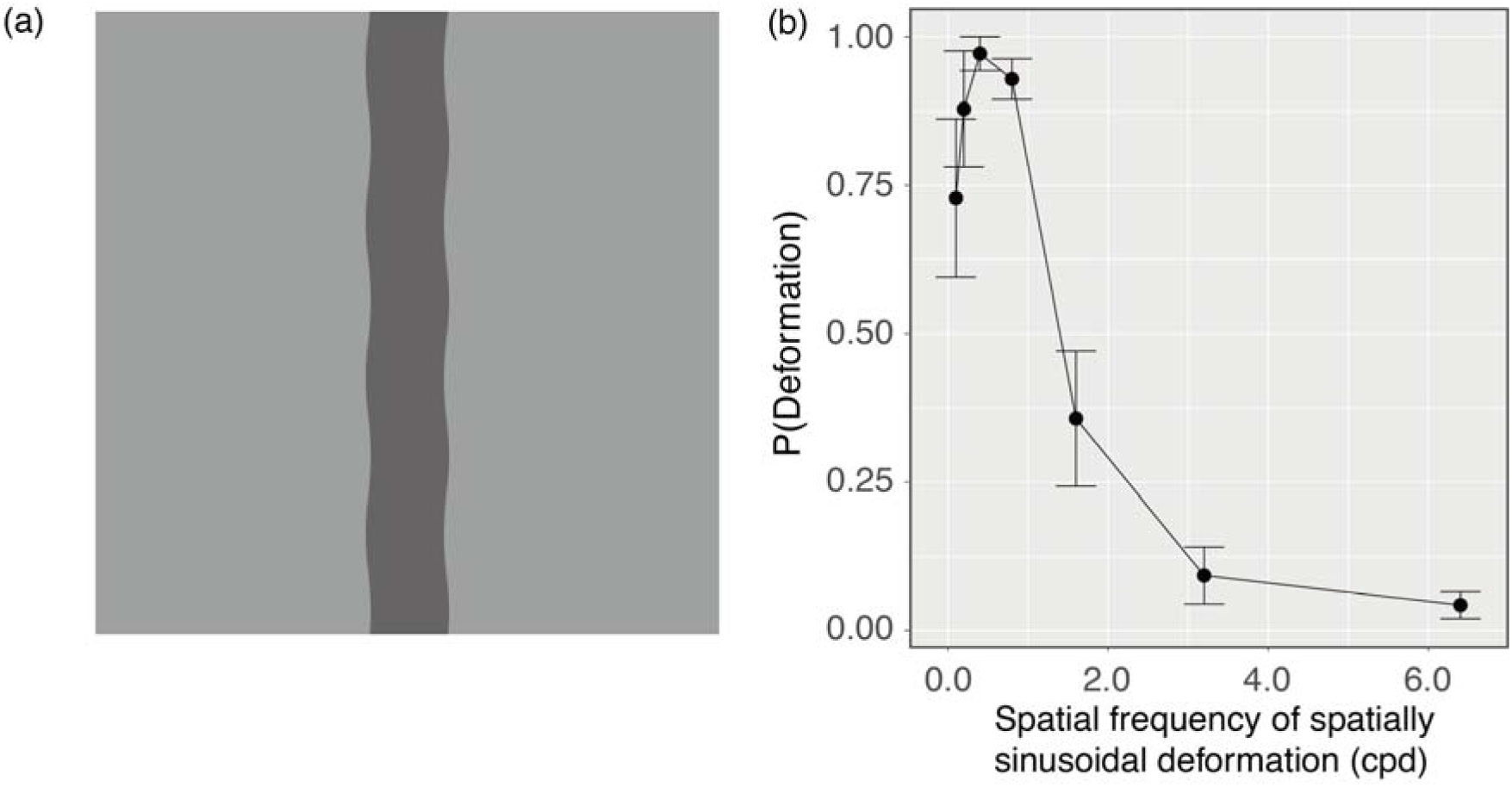
(a) A snapshot of a stimulus clip as used in Experiment 1 (Supplementary video 1). (b) Experiment 1 results. Error bars denote standard errors of the mean (N = 7).

**Figure 2.**
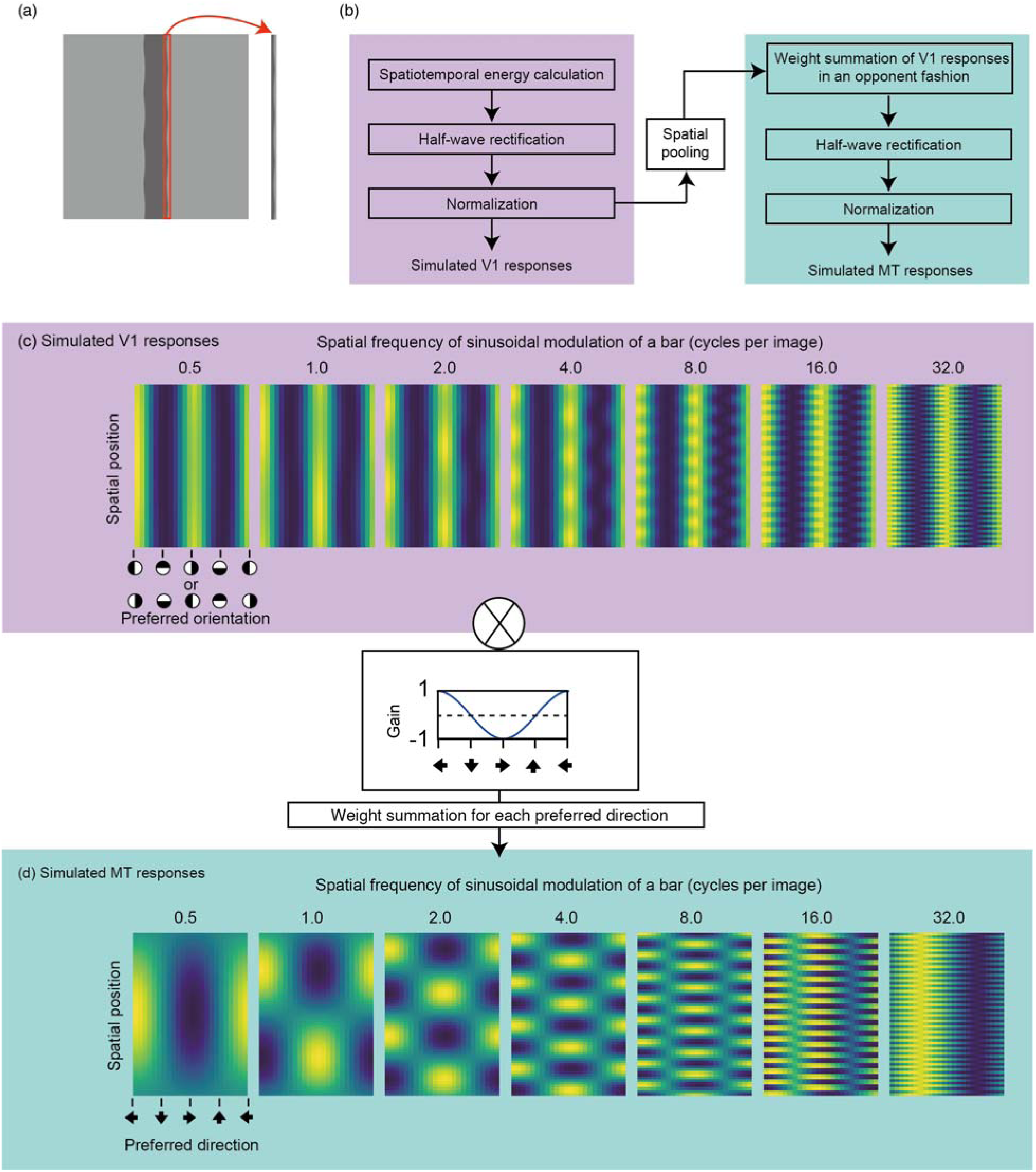
(a) Extracted area for simulation. The red-bound area was used to simulate the response of the direction-selective units. (b) A pipeline of our model. (c) Simulated spatiotemporal motion energy for the stimuli of Experiment 1. In this panel the range of each density plot is normalized between 0 and 1. Raw values were used for further analysis. (d) Simulated responses of direction-selective units for the stimuli of Experiment 1. In this panel the range of each density plot is normalized between 0 and 1. Raw values were used for further analysis.

**Figure 3.**
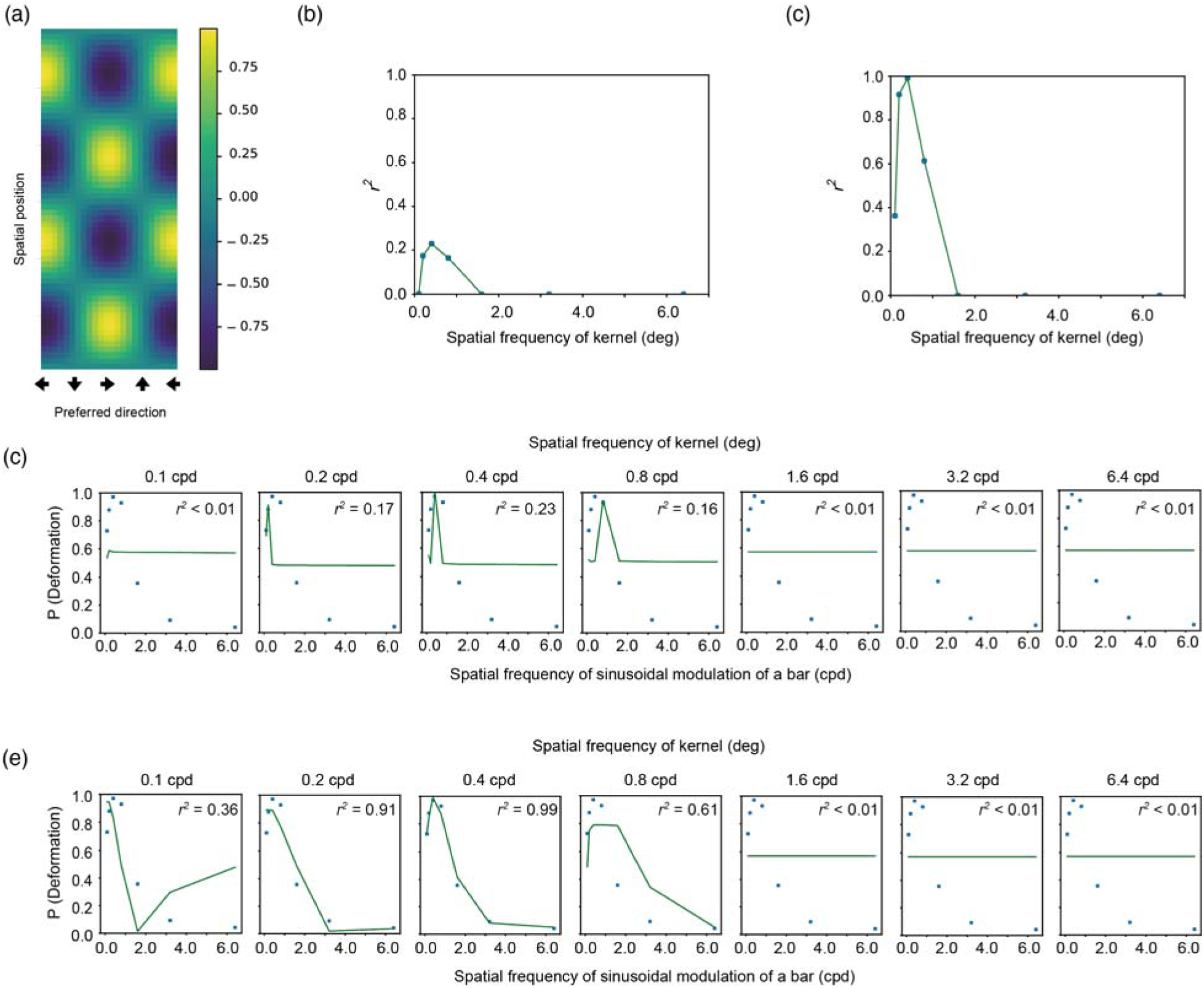
(a) An example of the kernel which was employed here. (b, c) The vertical axis denotes the coefficient determination (*r*^*2*^) for the fitting of an exponential function to the proportion of trials with deformation reports as a function of NCC, and the horizontal axis denotes the spatial frequency of modulation of a kernel. The panel b is for spatiotemporal motion energy and the panel c is for the response of direction-selective units. (d, e) The psychophysical data of deformation reports (markers) are jointly plotted with the fitted values (lines) as a function of the spatial frequency of sinusoidal deformation. The panel d is for spatiotemporal motion energy and the panel e is for the response of direction-selective units.

**Figure 4.**
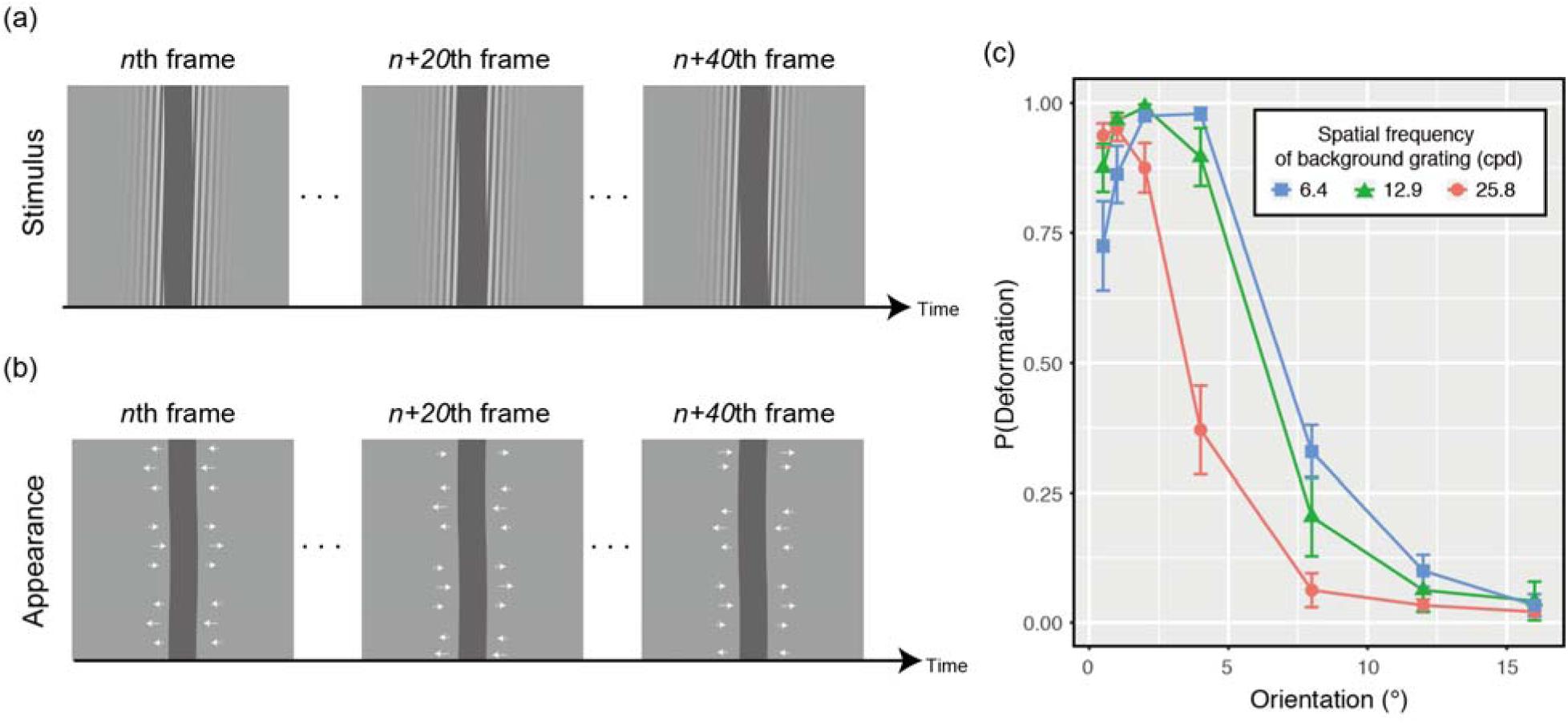
(a) Several snapshots of stimuli as used in Experiment 2. (b) Schematic explanations of the appearance of the deformation illusion of a bar on the basis of background drifting grating. (c) Proportions of deformation reports as a function of the orientation of background drifting grating for each spatial frequency condition of the background grating.

## Methods

### Experiment 1

#### Observers

7 people (5 females and 2 males) participated in this experiment. All observers in this study reported having normal or corrected-to-normal visual acuity. They were recruited from outside the laboratory and received payment for their participation. Ethical approval for this study was obtained from the ethics committee at Nippon Telegraph and Telephone Corporation (Approval number: H28-008 by NTT Communication Science Laboratories Ethical Committee). The experiments were conducted according to principles that have their origin in the Helsinki Declaration. Written, informed consent was obtained from all observers in this study.

#### Apparatus

Stimuli were presented on a 21-inch iMac (Apple Inc. USA) with a resolution of 1280 × 720 pixels and a refresh rate of 60 Hz. A colorimeter (Bm-5A, Topcon, Japan) was used to measure the luminance emitted from the display. A computer (iMac, Apple Inc., USA) controlled stimulus presentation, and data were collected with PsychoPy v1.83 (Peirce, 2007, 2009).

#### Stimuli

In the stimulus (Figure 1a and Supplementary Video 1), the edge of a vertical bar (0.6 deg width × 5.0 deg height) was horizontally deformed at one of the following 7 spatial frequencies (0.1, 0.2, 0.4, 0.8, 1.6, 3.2, and 6.4 cpd). We chose the range of spatial frequency of deformation to cover the range tested in the previous study (Nakayama & Silverman, 1988a). The amplitude was kept constant at 0.04 cpd. With each stimulus, upward or downward drifting was randomly given to the modulation. The modulation temporal frequency was 1 Hz. The luminance of the bar was randomly chosen as one of two levels (38 and 114 cd/m^2^). The luminance of the background was 76 cd/m^2^.

#### Procedure

Each observer was tested in a lit chamber. The observers sat 102 cm from the display. With each trial, a stimulus clip having a deforming bar was presented for 3 seconds. After the disappearance of the clip, a two-dimensional white noise pattern (with each cell subtending 0.16 deg × 0.16 deg) was presented until the observer’s response. The task of the observers was to judge whether a bar dynamically deformed or not. The judgment was delivered by pressing one of the assigned keys. Each observer had two sessions, each consisting of 7 spatial frequencies of the modulation × 10 repetitions. Within each session the order of trials was pseudo-randomized. Thus, each observer had 140 trials in total. It took approximately 20 minutes for each observer to complete all sessions.

### Experiment 2

#### Observers

12 people (10 females and 2 males) participated in this experiment. Their mean age was 38.2 (SD: 7.63). Although 7 of them also already participated in the previous experiment, none was aware of the specific purpose of the experiment because there was no preliminary explanation or debriefing provided.

#### Apparatus

Apparatus was identical to that used in Experiment 1.

#### Stimuli

As shown in Figure 4a (and Supplementary videos 2-4), a vertical bar (0.6 deg wide × 5.0 deg high) was presented in front of a drifting grating. For each stimulus, the luminance of the bar was randomly chosen from two levels (37 and 112 cd/m^2^). The orientation of the background grating was selected from the following 7 levels (0.5, 1, 2, 4, 8, 12, and 16°). The spatial frequency was selected from the following 3 levels (6.4, 12.9, and 25.8 cycles per degree). Drift temporal frequency was kept constant at 1 Hz. The drift direction was randomly determined. The luminance contrast of the grating was set at 0.75, and thus the luminance level of the grating ranged between 37 and 112 cd/m^2^. The drifting grating was windowed by a horizontal Gaussian envelope with a standard deviation of 0.62 deg.

#### Procedure

Procedure was identical to that in the previous experiment except for the following. With each trial, a stimulus clip having a static bar and drifting grating was presented for 3 seconds. After the disappearance of the clip, visual white noise (each cell subtending 0.16 deg × 0.16 deg) was presented until the observer’s response. The task of the observers was to judge whether the static bar dynamically deformed or not. Each observer had four sessions, each consisting of 3 spatial frequencies × 7 orientations × 5 repetitions. Within each session the order of trials was pseudo-randomized. Thus, each observer had 420 trials in total. It took 30–40 minutes for each observer to complete all four sessions.

### Experiment 3

#### Observers

12 people who had participated in Experiment 1 again participated in this experiment. Still, none was aware of the specific purpose of the experiment.

#### Apparatus

Apparatus was identical to that used in Experiment 1.

#### Stimuli

Stimuli were identical to those used in Experiment 1 except for the following. The background grating orientation was 1° or 16°, which respectively produced strong deformation and non-deformation responses in Experiment 1. The grating spatial frequency was kept constant at 12.9 cpd. As shown in Supplementary Video 5, the luminance of the bar was randomly chosen from the nine levels (0.0, 17.5, 37, 58, 76, 95, 112, 132, and 148 cd/m^2^ wherein 76 cd/m^2^ was the luminance of a neutral gray level).

#### Procedure

Procedure was identical to that used in Experiment 1 except for the following. Each observer had four sessions, each consisting of 2 levels of grating orientation × 9 luminance levels of the bar × 5 repetitions. Within each session the order of trials was pseudo-randomized. Thus, each observer performed 360 trials in total. It took approximately 20 minutes for each observer to complete all of both sessions.

### Simulation of MT responses

For the stimuli that were used in Experiments 1 and 2, we simulated the responses of direction-selective units on the basis of previous studies (Mante & Carandini, 2005; Nishimoto & Gallant, 2011; Perrone & Krauzlis, 2008; Perrone & Krauzlis, 2014; Simoncelli & Heeger, 1998). We extracted a stimulus area near the right vertical edge of the bar that was presented against a background (Figure 2a) and simulated the responses of the direction-selective units to the area.

#### Spatial parameters of spatiotemporal energy detection

In Experiment 1, we set the width of the extracted area for analysis at 0.08 deg because the amplitude of contour deformation applied to the bar was 0.04 deg. In Experiment 2, based on the spatial frequency of the background grating, we changed the width of the extracted area for analysis (0.16, 0.08, 0.04, deg for 6.4, 12.9, and 24.8 cpd conditions). The height of the extracted area was constant at 4.97 deg. The extracted area was first analyzed by a set of spatiotemporal filters. In Experiment 1, width, height, and spatial wavelength of the spatiotemporal filters were all 0.08 deg. In Experiment 2, width, height, and spatial wavelength of the spatiotemporal filters were also consistent with the width of the extracted area, that is, 0.16 deg, 0.08 deg, and 0.04 deg for 6.4, 12.9, and 24.8 cpd conditions, respectively. The filters with different phases (0π or 1.0π) were independently applied to stimuli and the outputs of the filters with different phases were later summed after the half-wave rectification and normalization described below. The number of filter orientations was 24 (i.e., 15° steps). The position of the filters did not spatially overlap. In Experiment 1, 64 filters covered the extracted area. In Experiment 2, 32, 64, and 128 filters covered the extracted area for 6.4, 12.9, and 24.8 cpd conditions, respectively.

#### Temporal parameters of spatiotemporal energy detection

In both experiments, the temporal size of the filter was 6 frames (for approximately 100 msec) and the temporal frequency was fixed at 0.4 Hz. The combination of temporal and spatial frequencies of the filter was optimized to the stimulus speed in the stimuli. We adopted the temporal properties because we wanted to extract a one-way modulation of the moiré pattern.

#### Rectification, normalization, and spatial pooling

The responses of the filters were half-wave rectified and normalized as reported in the previous study (Simoncelli & Heeger, 1998). In the calculation of normalization, as suggested by the previous study (Simoncelli & Heeger, 1998), the half-wave rectified output, which was multiplied by the maximum attainable response constant, was divided by the summed output of half-wave rectified responses across orientations and the semi-saturation constant. The normalized responses are considered as the spatiotemporal energy. The calculated motion energy is plotted in Figures 2c and 5b for Experiment 1 and Experiment 2, respectively. The normalized responses were spatially pooled among four adjacent filters, yielding 29, 61, and 125 responses. A Gaussian filter, which was centered on the pooling range, was applied to the pooling. The standard deviation of the Gaussian filter was 1.2.

#### Calculation of direction-selective responses

The pooled responses were filtered by a direction-tuned filter (Perrone & Krauzlis, 2008; 2014) that tuned to the motion direction using a cosine function. That is, at this level, the rectified and normalized outputs of the spatiotemporal energy filters were summed with weightings in an opponent fashion. The direction-tuned filters had 24 preferred directions (i.e., 15° steps). The filtered responses were half-wave rectified and then normalized as in a previous study (Simoncelli & Heeger, 1998). The constants were just identical to those as used in the previous study (Simoncelli & Heeger, 1998). This consequently yielded the responses of direction-selective units as functions of the preferred direction of units and spatial position, as shown in Figure 2d for Experiment 1 and Figure 5c for Experiment 2.

**Figure 5.**
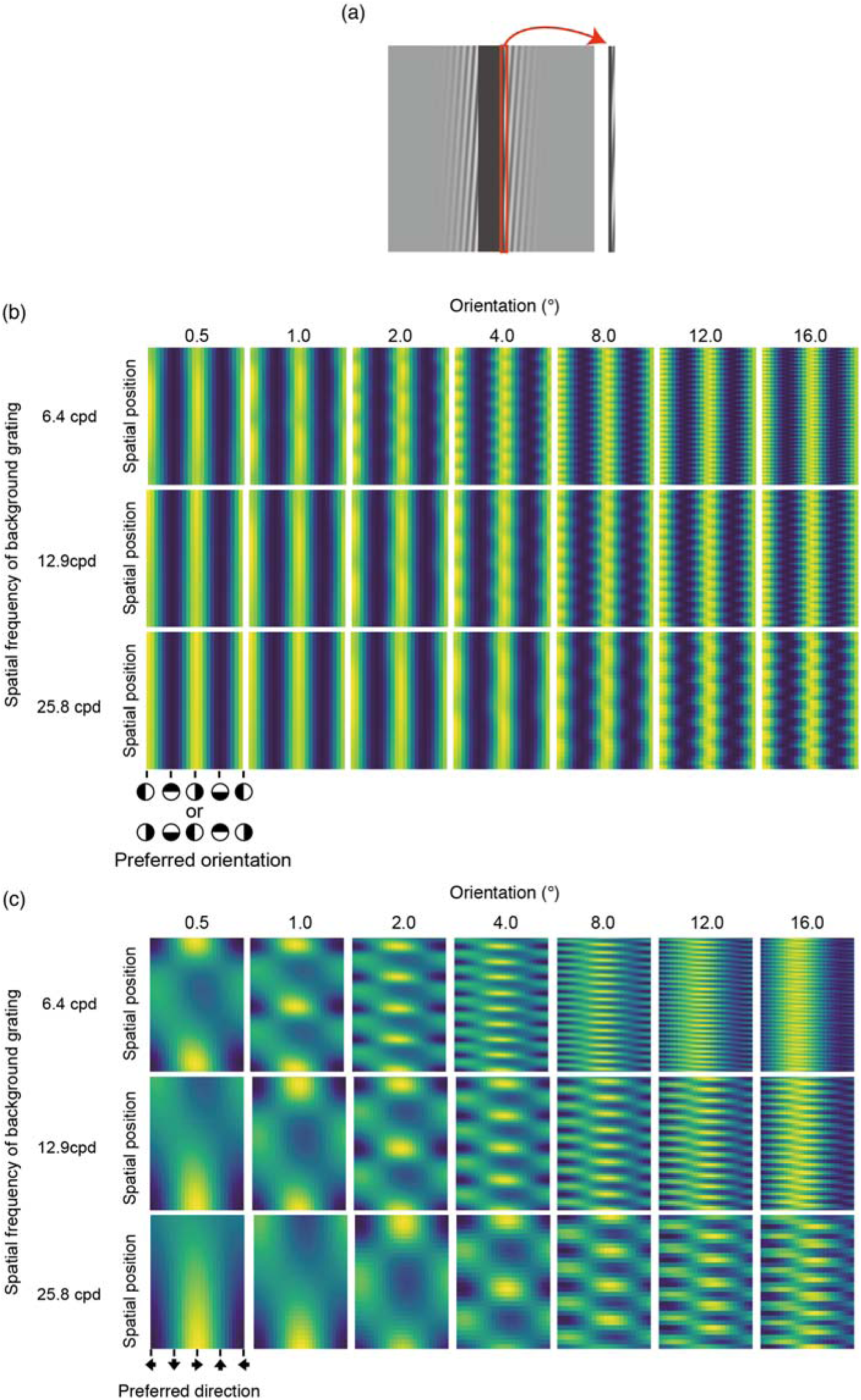
(a) Extracted area for simulation. (b) Simulated spatiotemporal motion energy for the stimuli of Experiment 1. In this panel, the range of each density plot is normalized between 0 and 1. Raw values were used for further analysis. (c) Simulated responses of direction-selective units for the stimuli of Experiment 1. In this panel, the range of each density plot is normalized between 0 and 1.Raw values were used for further analysis.

### Properties of units monitoring the spatial pattern of direction responses

The following analysis was conducted to calculate the normalized cross correlation (NCC) between spatiotemporal motion energy (or the response of direction-selective units) and the kernel of assumed higher-order units that are possibly sensitive to the spatial variation of spatiotemporal motion energy or the responses of direction-selective units. The kernel was defined by the product of the spatially sinusoidal pattern S and the directionally sinusoidal pattern D (Figure 3a). S was defined by the following formula,

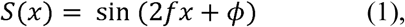

wherein *f* denotes spatial frequency, □ denotes phase, and x denotes spatial position. D was defined by the following formula,

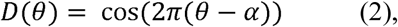

wherein *θ* denotes the motion direction and ranges from 0 to 2π, and α denotes the preferred direction of the kernel. Here, α was set to 0 deg (leftward direction) on the basis of the spatial pattern of the direction-selective units as shown in Figure 2d. Thus, a kernel K was defined as the product of S and D,

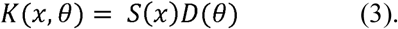

To see how the kernel (Figure 3a) matched the spatiotemporal motion energy (Figure 2c) or the response of direction-selective units (Figure 2d), we calculated the normalized cross-correlation (NCC) between them. The NCC was calculated by the following formula:

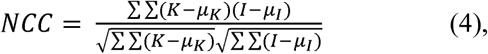

wherein *K* is the kernel, *I* is the spatiotemporal motion energy (or the pattern of direction-selective units), and *μ*_*K*_ and *μ*_*I*_ are the mean of *K* and *I*, respectively. This NCC is often called as the zero-mean NCC. The f of (1) took one of the following levels: 0.1, 0.2, 0.4, 0.8, 1.6, 3,2 and 6.4 cpd. The □ was tested in 32 steps (each step = 0.0625 π), and the maximum value among the 32 outcomes based on the 32 steps was considered to be the NCC of the kernel.

## Results and Discussion

### Experiment 1

The purpose of Experiment 1 was to specify psychophysical parameters to cause the perception of dynamic deformation and look into a possible relationship between the psychophysical data and the simulation data of neural responses such as V1 motion energy and MT pattern motion cells. Though a previous study (Nakayama & Silverman, 1988a) investigated amplitude thresholds at which dynamic deformation disappeared when a sinusoidally modulated line translated, no studies have directly examined which spatial frequencies of deformation produced the largest effect on deformation perception. Here we asked the observers to report whether the bar (which was actually deformed in the display) was seen as deforming, and examined the relationship between the observers’ responses and the spatial frequency of deformation.

Figure 1b shows the proportion of trials wherein the observers reported the bar as dynamically deforming. By using the proportion, we conducted a repeated-measures one-way ANOVA with the spatial frequency of modulation as a within-subject factor. The main effect was significant [*F*(6,36) = 38.358, *p* < .0001, *η*^*2*^_*p*_= 0.86]. Multiple comparison showed that the proportions in the 0.1, 0.2, 0.4, and 0.8 cycles per degree (cpd) conditions were significantly higher than the proportions in the 1.6, 3.2, and 6.4 cpd conditions (*p* < .05). The results showed that the lower spatial frequency of deformation contributed to the perception of dynamic deformation more strongly than the higher spatial frequency of deformation, consistent with the previous study (Nakayama & Silverman, 1988a) showing that the amplitude thresholds at which dynamic deformation was abolished increased for lower spatial frequencies of physical deformation of a line. When the spatial frequency of deformation was high, some observers reported that they saw a rigid translation of wavy patterns along the edge of a bar; this was also consistent with the previous study (Nakayama & Silverman, 1988a).

### Simulation of MT responses

To specify the mechanism for seeing dynamic deformation in our stimuli, we decided to check the relationship between the perception of dynamic deformation and the simulated responses of units at the V1 and MT levels. On the basis of the previous literature (Mante & Carandini, 2005; Nishimoto & Gallant, 2011; Perrone & Krauzlis, 2008; Perrone & Krauzlis, 2014; Simoncelli & Heeger, 1998), we constructed the simulation by employing a standard procedure with the following steps (see also Figure 2b and Methods for details):

1. Convolving stimuli with spatiotemporal filters to get spatiotemporal motion energy.
2. Half-wave rectification.
3. Divisive normalization. At this stage, spatiotemporal energy of stimuli was obtained.
4. Spatial pooling with a spatially gaussian window.
5. Weighted summation of spatiotemporal energy in an opponent fashion.
6. Half-wave rectification.
7. Divisive normalization. At this stage, the response of a direction-selective unit was obtained.

Figure 2c shows the spatiotemporal motion energy as functions of the preferred spatiotemporal orientation and spatial position, for each condition of the spatial frequency of sinusoidal deformation. Figure 2d shows the responses of direction-selective units as functions of the preferred direction of the units and spatial position, for each condition of the spatial frequency of sinusoidal deformation. For both spatiotemporal energy and the responses of direction-selective units, the spatial pattern got finer as the spatial frequency of deformation increased. The spatiotemporal motion energy was high at the vertical orientations consistently across space. On the other hand, the response of direction-selective units was high at the leftward and rightward but in a spatially alternating manner.

### Properties of units monitoring the spatial pattern of direction responses

Based on the simulation, we next examined whether the psychophysical results could be well accounted for by spatiotemporal motion energy and/or the responses of direction-selective units. The brain would determine whether the input signal came from deformation or not, based on the output of the higher-order unit that is sensitive to the spatial pattern of spatiotemporal motion energy and/or the responses of direction-selective units. To test this prediction, we conducted a pattern-matching analysis using a kernel as shown in Figure 3a. From the results of the simulation of the spatiotemporal motion energy (Figure 2c) and direction-selective units (Figure 2d), it was plausible to assume that the higher-order units could tune to spatially sinusoidal modulation of motion direction as shown in Figure 3a. We call the template pattern having the spatially sinusoidal modulation “a kernel”. To assess the similarity between the kernel and each spatial pattern of spatiotemporal motion energy and the spatial pattern of the response of direction-selective unit, manipulating the spatial frequency of the modulation in the kernel, we calculated the normalized cross-correlation (NCC) between the kernel and the spatial pattern of the spatiotemporal motion energy or the responses of direction-selective units. We assumed that the calculated NCC could account for the psychophysical data for deformation perception. It was thus expected that the NCC would well explain the psychophysical data when the kernel had a high correlation with the simulated neural responses that effectively contribute to the deformation perception. We calculated NCCs between a kernel with one of the pre-determined spatial frequency of modulation and the spatial pattern of the spatiotemporal motion energy (Figures 2c) and/or the simulated responses of direction-selective units (Figure 2d).

Next, we fitted an exponential function to the psychophysical data as a function of the NCCs. The coefficient of determination (*r*^*2*^) of the fitting is plotted in Figure 3b for the spatiotemporal motion energy and in Figure 3c for the responses of direction-selective units. The psychophysical data (markers) and fitted values (lines) are jointly plotted in Figure 3d for the spatiotemporal motion energy and in Figure 3e for the response of direction-selective units as a function of the spatial frequency of sinusoidal deformation. For both the spatiotemporal motion energy and the responses of direction-selective units, the lower bands of the spatial frequency of the modulation in the kernel showed the highest coefficient of determination. The highest coefficient of determination implies that for the brain the information at the lower spatial frequency range reliably contributes to the estimation of dynamic deformation. Moreover, the coefficient of determination was higher for the response of direction-selective units than for the spatiotemporal motion energy.

The results suggest that the perception of bar deformation is likely mediated by the higher-order unit monitoring the spatial pattern of the responses of the direction-selective units. Moreover, the observers’ reports for deformation perception seem tuned to the lower spatial frequency of the responses of direction-selective units. The results are well consistent with the previous study showing that the spatial frequency of image deformation determines the appearance of image deformations (Kawabe, Maruya, & Nishida, 2015). The higher coefficient of determination for the response of direction-selective units than the spatiotemporal motion energy indicates that deformation perception is based on the direction responses at the MT area rather than the spatiotemporal motion energy at the V1 area.

The simulated response of the V1 cell was spatially sinusoidal but a little bit noisy. The noisiness of the response might come from that the orientation of a stimulus edge was near-vertical. Thus, there was a possibility that most of the V1 cells that were sensitive to the vertical orientation captured the signals of the edges, making the baseline of their activity non-zero along the edge. The high baseline possibly attenuated the sinusoidal pattern of the V1 responses. On the other hand, the response of direction-selective units does not depend strongly on the stimulus orientation because the MT cells solve the aperture problem. The solution of the aperture problem possibly made the simulated response of the direction-selective units less noisy than the simulated response of the V1 cells.

When the spatial frequency of the sinusoidal modulation of a bar was high, the simulated responses of direction-selective units showed preferences for downward motion (Figure 2d). This pattern of responses is consistent with the psychophysical data in a previous study (Nakayama & Silverman, 1988a) which showed that the sinusoidal modulation of a line perceptually resulted in unidirectional translation when it had a high spatial frequency of modulation. The results of our simulations indicate that the model we employed could precisely capture the properties of human motion perception with stimuli containing dynamic deformation.

### Experiment 2

The purpose of this experiment was to confirm whether illusory deformation perception that is induced by Moiré pattern (Figures 4a and 4b) could also be explained by the activities of higher-order units that are tuned to the spatial pattern of responses of direction-selective units.

We calculated the proportion of trials in which the observer reported the dynamic deformation of the bar and plotted them as a function of grating orientation for each spatial frequency condition in Figure 4c. We conducted a two-way repeated-measures ANOVA with grating orientation and grating spatial frequency as within-subject factors. The main effect of the grating spatial frequency was significant [*F*(2,22) = 27.440, *p* < .0001, *η*^*2*^_*p*_= 0.72]. The main effect of the grating orientation was also significant [*F*(2,22) = 255.772, *p* < .0001, *η*^*2*^_*p*_= 0.96]. Interaction between the two factors was significant [*F*(12,132) = 15.362, *p* < .0001, *η*^*2*^_*p*_= 0.58). The results showed that the illusory deformation occurred when the background drifting grating had smaller tilts away from vertical. At the same time, there was a significant interaction between orientation and spatial frequency of the background grifting grating. The significant interaction possibly comes from the peak shift of the deformation reports which occurred depending on the spatial frequency of background grating. As the spatial frequency of background grating decreased, the peak of deformation reports occurred at the shallower orientation of the grating.

We surmised that the peak shift might come from the change in the activity of the higher-order units that were sensitive to the spatial pattern of the responses of direction-selective units, and hence decided to simulate the responses of direction-selective units for the stimuli as used in this experiment. In a similar way to the previous analysis, by manipulating the spatial frequency of the kernel (Figure 3a) we calculated the NCC for each stimulus and fitted an exponential function to the psychophysical data as a function of the NCCs. The coefficients of determination (*r*^*2*^) of the fitting are shown in Figure 6a for the spatiotemporal motion energy and in Figure 6b for the response of direction-selective units. As in the previous analysis, *r*^*2*^ peaked at the lower spatial frequency bands of the spatial pattern of direction responses. The psychophysical data and fitted values are jointly plotted in Figure 6c for the spatiotemporal motion energy and in Figure 6d for the response of direction-selective units as a function of the spatial frequency of sinusoidal modulation. Similar to the results of Experiment 1, the coefficient of determination was higher for the response of direction-selective units than for the spatiotemporal motion energy.

**Figure 6.**
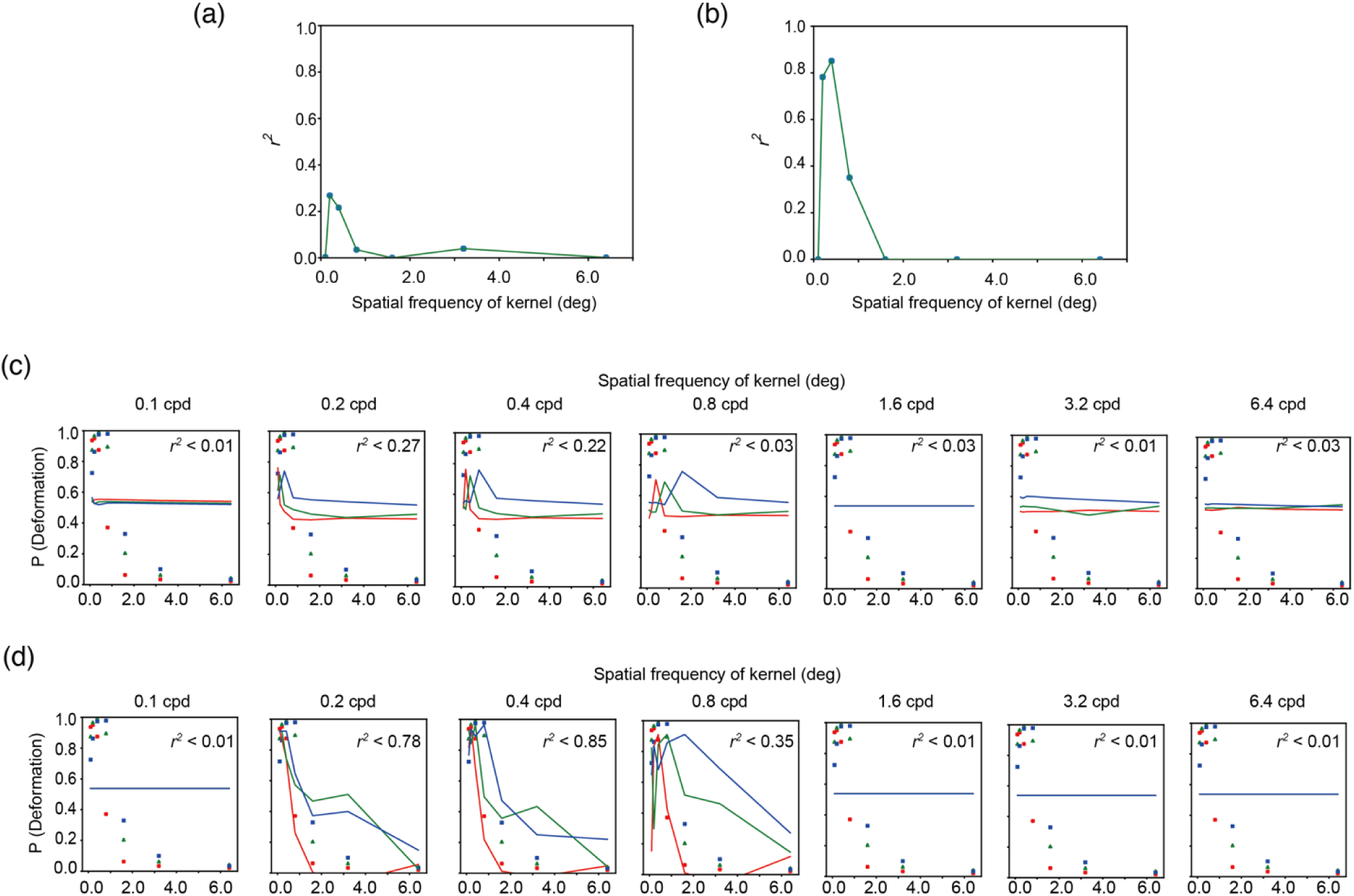
(a, b) The vertical axis denotes the coefficient determination (*r*^*2*^) for the fitting of an exponential function to the proportion of trials with deformation reports as a function of NCC, and the horizontal axis denotes the spatial frequency of modulation of a kernel. The panel a is for spatiotemporal motion energy and the panel b is for the response of direction-selective units. (c, d) The psychophysical data of deformation reports (markers) are jointly plotted with the fitted values (lines) as a function of the spatial frequency of sinusoidal deformation. The panel c is for spatiotemporal motion energy and the panel d is for the response of direction-selective units.

The results indicate that similar to the stimuli that contained the actual deformation of a bar, the brain uses the output of the higher-order units that are sensitive to the spatial pattern of responses of direction-selective units in determining illusory deformation perception. Moreover, consistent with the results of Experiment 1, the response of direction-selective units at the MT area possibly more strongly contributes to the illusory deformation perception than the spatiotemporal motion energy.

### Experiment 3

The purpose of this experiment was to additionally confirm whether the deformation perception could be obtained on the basis of the contrast-based Moiré pattern. In Experiment 2, we used dark and bright bars against a background drifting grating, so the Moiré pattern generated between the bar and the background grating was always defined by luminance. By testing the condition with the bar luminance at a neutral gray (Supplementary video 5), we investigated whether the contrast-based Moiré pattern also contributed to the perception of dynamic deformation. If the deformation perception was operated by the mechanism that had a function equivalent to the direction-selective units, because the unit was assumed to have selectivity to luminance, no strong influence of contrast-based Moiré pattern would be observed.

In Figure 7, the proportion of trials with reports of bar deformation is plotted for each background orientation condition as a function of the luminance of the bar. As in Experiment 1, the deformation perception was more often reported with a grating orientation of 1° than 16°. Interestingly, the proportion suddenly dropped when the luminance of the bar was set at the level of neutral gray. By using the proportion, we conducted a two-way repeated-measures ANOVA with the bar luminance and background grating orientation as within-subject factors. The main effect of the bar luminance was significant [*F*(8,88) = 16.36, *p* < .0001, *η*^*2*^_*p*_= 0.60]. The main effect of the background grating orientation was also significant [*F*(1,11) = 1463.35, *p* < .0001, *η*^*2*^_*p*_= 0.99]. Interaction between the two factors was also significant [*F*(8,88) = 17.763, *p* < .0001, *η*^*2*^_*p*_= 0.62]. The simple main effect based on the significant interaction showed that the proportion for 76 cd/m^2^ was significantly lower than the proportions for other luminance conditions when the background orientation was 1° (*p* < .05). The results showed that the deformation perception was attenuated when the bar luminance was neutral gray, suggesting that the mechanism for the deformation perception in our illusion is most sensitive when there is a difference in overall luminance between a bar and its background. Moreover, the illusory deformation was still reported even when the luminance of the bar itself was outside of the luminance contrast range of the background grating. The results suggest that the deformation illusion triggered by moiré patterns occurs with a flexible relationship between the bar and the background grating.

**Figure 7.**
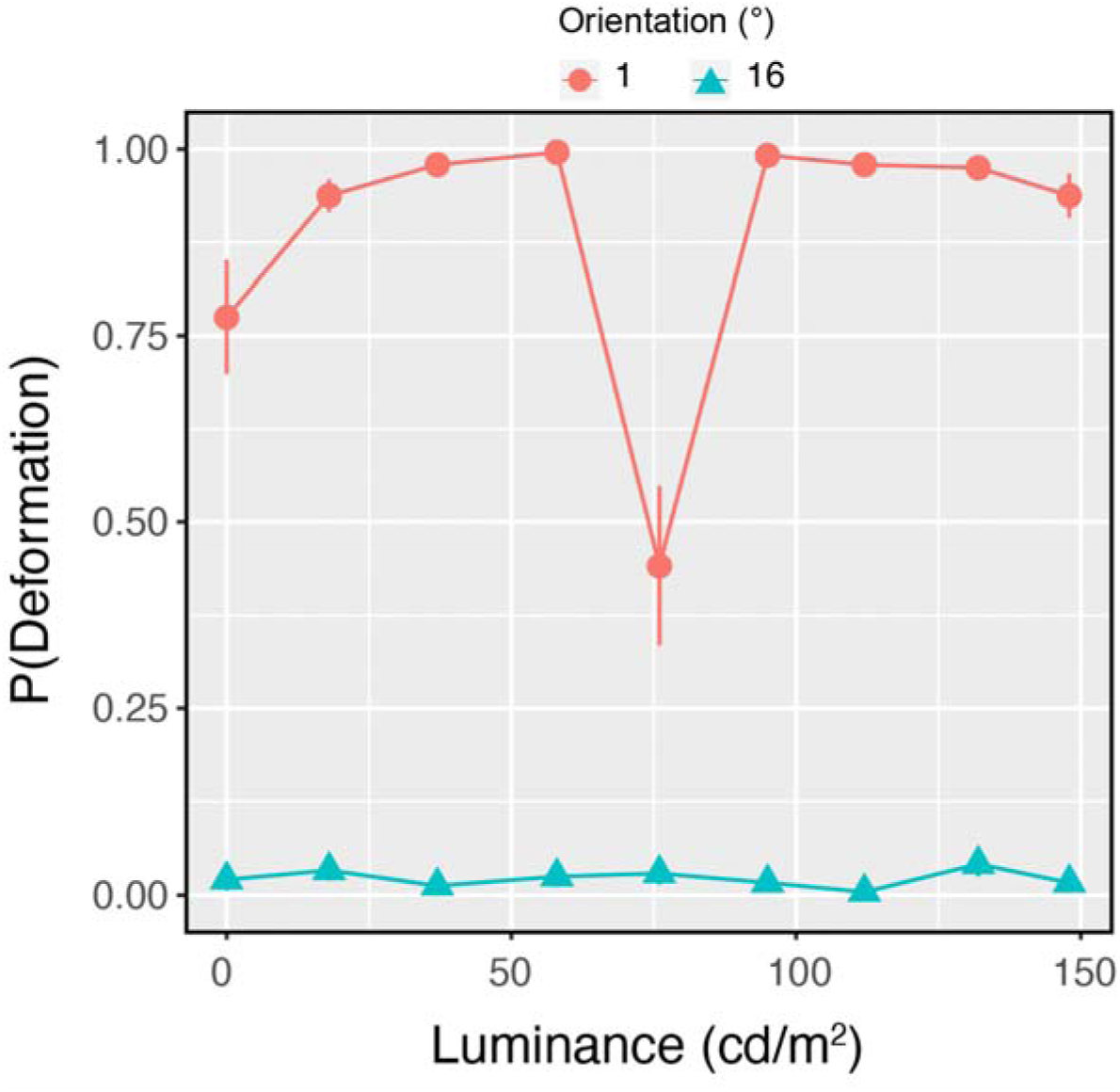
Experiment 3 results. Proportions of deformation reports as a function of the luminance of a bar in stimuli. Error bars denote ±1 standard errors of the mean for each of the orientation conditions of background drifting grating.

## General Discussion

This study investigated the mechanism responsible for deformation perception by human observers. Our data suggest that the brain determines deformation perception on the basis of the spatial pattern of the responses of the direction-selective units. Moreover, it was shown that the mechanism underlying the deformation perception was more selective to luminance-defined than contrast-defined features.

Because the spatiotemporal motion energy did not well explain the deformation perception in our stimuli, we believe that the deformation perception is not based on a series of static deformations. Rather, it seems plausible to assume that motion mechanisms solving the aperture problem produce the kind of dynamic deformations, as reported in the previous study (Nakayama & Silverman, 1988a). Besides, in both Experiments 1 and 2 of this study, the response of the direction-selective units showed the preference for downward motion when the spatial frequency of the sinusoidal deformation of a bar was high, while the spatiotemporal motion energy did not show such preference for spatiotemporal orientation involving downward motion. The downward preference is also explained in terms of solving the aperture problem (Nakayama & Silverman, 1988a). As such, rather than the spatiotemporal motion energy, the response of direction-selective units at the MT area better characterizes the perception of dynamic deformation.

Previous studies have reported that the human visual system is sensitive to direction-defined stripes (van Doorn & Koenderink, 1982a, 1982b) and/or direction-defined gratings (Nakayama et al., 1985). Does the mechanism responsible for motion-defined structures also mediate the perception of deformation? Previous studies have consistently shown that the mechanism for the detection of a motion-defined structure is low-spatial-frequency selective. On the other hand, as shown in Figures 3c and Figure 4c, the mechanism for deformation perception may have band-pass properties though relatively lower bands possibly contribute to the deformation perception. We, therefore, suggest that the mechanism for deformation perception is not simply equivalent to the mechanism for detecting motion-defined structures, though it is possible that some processing procedures are shared between them.

What is the possible neural mechanism for seeing deformation? It is well known that velocities are processed in the MT area. In general, velocities are captured via a large receptive field and hence processed globally (Dubner & Zeki, 1971; Newsome et al., 1985). On the other hand, some studies have reported that an identical receptive field of the MT area could locally respond to independent velocities (Majaj et al., 2007; Perrone & Krauzlis, 2008). The kind of local velocity extraction may mediate the perception of deformation, though further clarification is necessary to evaluate this possibility since the spatial properties of the local velocity extraction in the MT area are still an open issue. Structure from motion, which is occasionally involved with a complex motion structure, is also processed in the MT area (Bradley et al., 1998; Grunewald et al., 2002). There is thus a possibility that the MT area is responsible for deformation perception, which is also involved with complex motion structure.

In this study, we did not closely investigate the role of speed in the perception of dynamic deformation. Specifically, we assessed only direction parameters in the computation model, while the model proposed in previous studies (Simoncelli & Heeger, 1998) could assess both direction and speed. This was because the most revealing speed of image motion signals in our stimuli could easily anticipated for both illusory and real bar deformations, and so it was possible for us to determine the optimal speed parameter of the model in advance. On the other hand, it is known that local motion speeds determine the appearance of image deformation (Kawabe, 2018). Manipulating the speed of deformation as well as checking the speed parameters in the MT model need to be tested in future investigations.

How is the illusory deformation in our stimuli related to the footstep/inchworm illusion? In the footstep illusion (Anstis, 2001, 2004; Howe et al., 2006), an object translating at a constant velocity apparently changes its speed depending on the luminance contrast between the object and a black-white stripe background. The illusion occurs even when the contrast between the object and the background is defined by second-order features (Kitaoka & Anstis, 2015). In the inchworm illusion, a similar sort of the contrast effect on apparent speed produces the extension and contraction of the object along its motion trajectory (Anstis, 2001). The footstep/inchworm illusion is, at a glance, similar to the illusory deformation in our stimuli in terms of that the background stripe produces the change in motion appearance of a foreground object. However, there is a critical difference in the appearance between them. In the footstep illusion, the object apparently stops or reduces its speed when the contrast between the moving object and its background is low. On the other hand, in the illusion the present study reported, the static object apparently deforms when the contrast between the object and the background grating is low. That is, observers see the illusory deformation when the direction signals are produced at the intersection between, for example, a static black object and the dark part of the drifting grating. In this respect, different mechanisms basically underlie between the footstep/inchworm illusion and the illusory deformation in the present study. The footstep illusion is related to a contrast-based speed illusion and the illusory deformation in this study is related to the direction-based illusion based on the Moiré pattern. According to the previous study (Anstis, 2001), the footstep illusion can also produce the two-dimensional direction illusion when the background is replaced with the two-dimensional checker-board like pattern. Anstis (2001) suggested that the apparent directional change could be predicted from the vector averaging of local motion. It is assumed that the kind of vector averaging is mediated by the processing in the MT area (Simoncelli & Heeger, 1998). Thus, the mechanism for determining direction may be common between the footstep/inchworm illusion and the illusory deformation in our study.

We acknowledge that the results of Experiment 3 are explained in terms of the act of spatiotemporal filters at the first stage of our model. To detect contrast-defined features, we need to assume nonlinear processing before evaluating the spatiotemporal structure of contrast modulation. The spatiotemporal filter does not, in general, follow a nonlinear processing as like a rectification and hence it is impossible for the simple spatiotemporal filters to detect the contrast-defined structures. In Experiment 3, the deformation perception was attenuated when the overall luminance of the background was equivalent to a bar. The results are consistent with the interpretation that the illusory deformation perception occurs when the spatiotemporal filters that are sensitive to luminance structures properly detect the spatiotemporal structures that eventually causes the activation of the direction selective units producing the lower spatial frequency pattern. On the other hand, we would like to emphasize that after the detection of spatiotemporal luminance structure the brain solves an aperture problem to determine global motion directions when deformation is dynamic. The solution to the aperture problem is mediated by the filters at MT. Thus, we need to assume the two-stage processing to entirely describe the dynamic deformation perception in our stimuli.

In this study, we focused on shearing deformation but not on compressive deformation. Both shearing, and compressive deformations exist in the image deformation of natural materials such as the flow of a transparent liquid. Showing that the sensitivity to the compressive deformation was higher than the sensitivity to the shearing deformation, Nakayama et al. (1985) has proposed that different mechanisms mediate the compressive deformation from the shearing deformation. On the other hand, just how shearing and compressive deformation are processed and interact with each other in the visual system remains an open question. Psychophysical and computational investigation of simultaneous shearing and compressive deformations will lead to further understanding of how the visual system detects and interprets image deformation in natural scenes.

